# SARS-CoV-2 BA.4 infection triggers more cross-reactive neutralizing antibodies than BA.1

**DOI:** 10.1101/2022.07.14.500039

**Authors:** Simone I. Richardson, Thopisang Motlou, Mieke A. van der Mescht, Bronwen E. Lambson, Josie Everatt, Daniel Gyamfi Amoako, Anne von Gottberg, Nicole Wolter, Zelda de Beer, Talita Roma de Villiers, Annie Bodenstein, Gretha van den Berg, Theresa M. Rossouw, Michael T. Boswell, Veronica Ueckermann, Jinal N. Bhiman, Penny L. Moore

## Abstract

SARS-CoV-2 variants of concern (VOCs) differentially trigger neutralizing antibodies with variable cross-neutralizing capacity. Here we show that unlike SARS-CoV-2 Omicron BA.1, which triggered neutralizing antibodies with limited cross-reactivity, BA.4/5 infection triggers highly cross-reactive neutralizing antibodies. Cross-reactivity was observed both in the absence of prior vaccination and also in breakthrough infections following vaccination. This suggests that next-generation vaccines incorporating BA.4, which is spreading globally, might result in enhanced neutralization breadth.

## Main text

SARS-CoV-2 variants of concern (VOCs) which contain mutations within the spike gene have been associated with viral escape from neutralization^1-4^, and thus reduced vaccine efficacy^5^. Most recently, these VOCs include Omicron BA.1, which contains >30 mutations in the spike region, resulting in further reduced neutralization titers^6^. Omicron has since evolved into several sub-lineages including BA.2, BA.2.12.1, BA.4 and BA.5^7^. The BA.4 and BA.5 sub-lineages, which share the same dominant spike mutations but differ from one another in non-structural proteins as well as the nucleocapsid and matrix genes, drove the fifth wave of infection in South Africa, and have subsequently been detected in more than 30 other countries^8^. Within spike, BA.4 and BA.5 are genetically similar to BA.2 but contain two the 69-70 deletion, additional substitutions in the receptor binding domain (RBD), namely L452R, F486V and the R493 reversion to wild-type. Therefore, compared to BA.1 and BA.2, BA.4 has shown increased neutralization resistance to vaccinee and convalescent sera, and monoclonal antibodies^4,9,10^.

In addition to conferring variable escape, mutations in spike also result in the elicitation of qualitatively different responses after infection. We and others have shown that each SARS-CoV-2 variant triggers different profiles of neutralizing antibodies (nAbs)^2,3,11-14^. For example, the Beta variant triggered humoral responses with increased cross-reactivity, and consequently became a focus for the development of second-generation vaccines^14-21^. In contrast, Omicron BA.1 triggered more strain-specific nAbs with limited cross-reactivity in individuals without prior infection. This likely reflects its significant divergence from the wild-type variant and suggests that BA.1 may not be an optimal insert for next generation vaccines^11,12^. Here, we assessed the cross-reactivity of nAbs triggered by BA.4 infection in previously vaccinated and unvaccinated individuals and compared the degree of cross-reactivity with sera from previous waves of infection in South Africa, caused by D614G, Beta, Delta or BA.1.

Plasma from individuals infected during the fifth wave of the COVID-19 pandemic in South Africa were used to assess neutralization cross-reactivity. Samples were obtained from 35 hospitalized individuals from the Tshwane District Hospital recruited between 9 and 22 May 2022, at a median of two days (range 0 to 9 days) after a positive PCR test (Table S1). During this period, national sequencing showed that Omicron BA.4/5 was responsible for >90% of genomes^8^. Of these participants, 30 had matched nasal swabs available. Whole genome sequencing data with more than 50% genome coverage was generated for 21/30 samples and 67% (14/21) of these could be assigned a clade and lineage. Clade and lineage assignments reflected that 79% (11/14) and 21% (3/14) of these Omicron sequences were BA.4 and BA.5 respectively. Thirteen individuals had previously been vaccinated with either one dose of Ad26.CoV2.S (n = 5) or two doses of BNT162b2 (n = 8) at least two months (range 56 to 163 days) prior to infection (Table S1). Twenty-two individuals were unvaccinated and had no history of previous symptomatic COVID-19 infection (Table S1).

Plasma from unvaccinated individuals were screened against the sequence matched BA.4/5 pseudovirus and 41% (9/22) showed no detectable neutralization (Figure S1). We note that this is a higher proportion of non-responders than we previously observed in the fourth wave of infections caused by BA.1 (20% non-responders)^12^. Of note, 8/9 unvaccinated non-responders were hospitalised for other reasons and were incidental SARS-CoV-2 detections, with mild or asymptomatic infections. In contrast, a lower fraction of the twelve vaccinated individuals were non-responders (n = 3), and all had previously received the Ad26.CoV2.S vaccine.

We used a lentiviral pseudovirus neutralization assay to assess cross-reactivity to the ancestral D614G, Beta, Delta, and Omicron BA.1, BA.2 and BA.4, excluding non-responders (individuals who failed to mount autologous responses to BA.4/5 (non-responders). In unvaccinated individuals, we observed moderate titers against the matched BA.4/5 spike (geometric mean titer (GMT) of 1,047 in those with detectable neutralizing titers). These moderate titers are consistent with the mild/asymptomatic infections these individuals had (Table S1). Against VOCs, the fold reduction in neutralization titers was variable, with titers against D614G, Beta, and BA.2 not significantly lower than BA.4/5 (GMT of 904, 859 and 884 respectively). Tiers against BA.1 were 2-fold lower (GMT of 557) and those against Delta were significantly lower, with a GMT of 258.

We and others have previously shown that breakthrough infection (BTI) following vaccination results in significantly boosted titers^12,22,23^. Here, in nine previously vaccinated individuals with BTI, we observed significantly boosted titers against BA.4, with a GMT of 1984. Titers were slightly enhanced against D614G consistent with prior exposure to the vaccine insert (Wuhan-1). Against other VOCs, we observed a slight reduction in titers, none of which were significant, with GMTs of 2972 for Beta to 4244 for BA.2 (Figure 2). Overall, therefore in previously vaccinated BTIs, neutralizing responses triggered by BA.4 BTI triggered high titers of cross-reactive antibodies.

**Figure 1:**
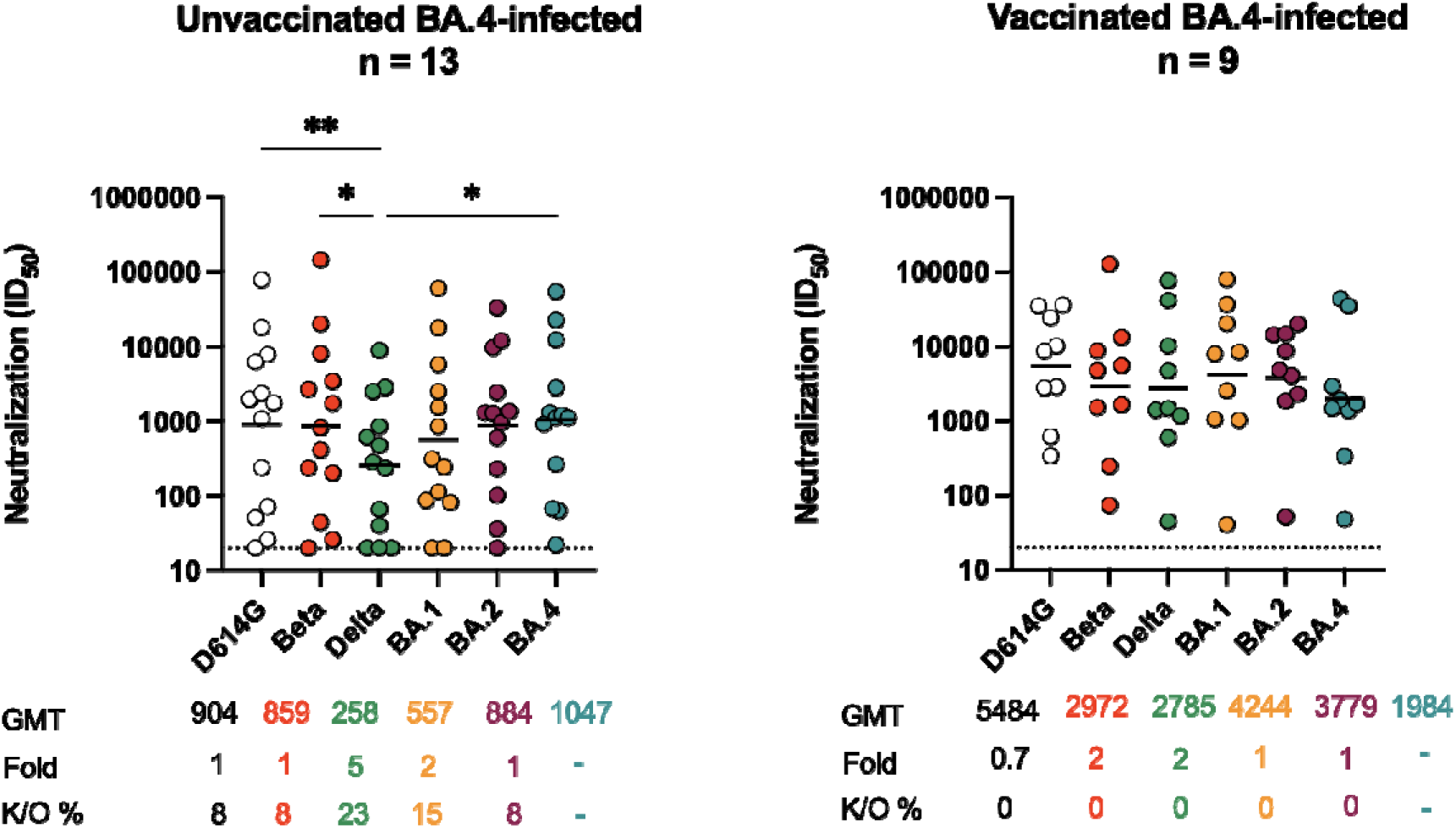
Omicron BA.4 triggers cross-variant neutralizing antibodies which are broadened by vaccination. Neutralization titer (ID_50_) of Omicron BA.4-infected plasma against D614G, Beta, Delta, Omicron BA.1, BA.2 and BA.4 pseudoviruses shown for (A) unvaccinated individuals (n=13) or (B) individuals vaccinated with either one dose of Ad26.CoV.2S or two doses of BNT162b2 (n=9). Lines indicate geometric mean titer (GMT) also represented below the plot with fold decrease and knock-out (K/O) of activity for other variants as a percentage relative to Omicron BA.4. Dotted lines indicate the limit of detection of the assay. Statistical significance across variants is shown by Friedman test with Dunns correction. All data are representative of two independent experiments.

**Figure 2:**
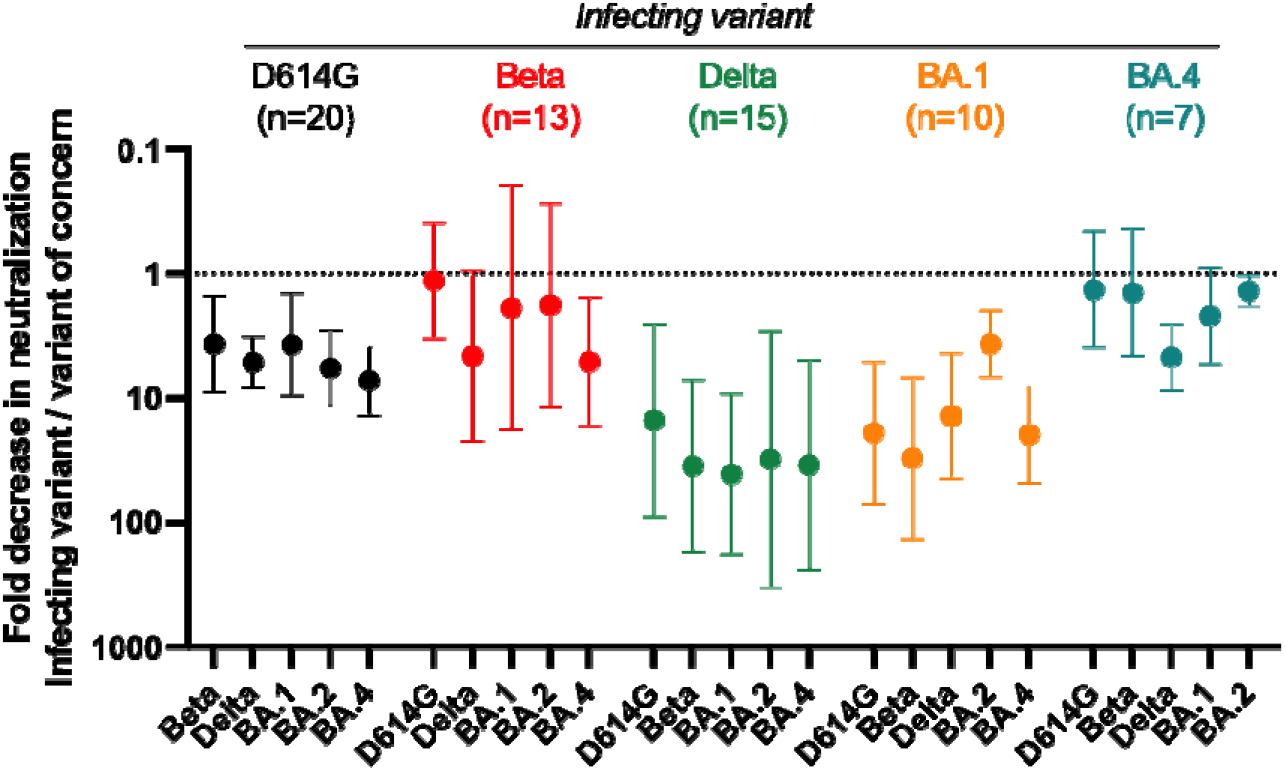
Comparison of cross-reactivity of antibodies elicited by infection D614G, Beta, Delta, BA.1 and BA.4 infection in previously unvaccinated individuals. Fold decrease in neutralization for each VOC represented as a ratio of the titer to the infecting variant (D614G, Beta, Delta, BA.1 and BA.4/5) over titer against unmatched VOCs. Shown are geometric means of the ratios, with error bars denoting the 95% CI. All data are representative of two independent experiments.

To compare the breadth of BA.4/5 triggered responses in unvaccinated individuals, we performed a meta-analysis of previously described responses in infections caused by each of the waves in South Africa (D614G, Beta, Delta, BA.1 and BA.4/5)^12-15^. To assess cross-reactivity, we measured the ratio of the neutralization titer against the infecting virus over the titer of the same plasma sample against each VOC, with values <1 representing loss of activity against variants (Figure 2). We show that while Beta and BA.4/5 triggered responses that are highly cross-reactive, the Delta and BA.1 variants triggered responses with less cross-reactivity, with 10-100 fold reductions in titer against many VOCs (Figure 2).

Overall, these data suggest that, in the absence of prior vaccination, unlike BA.1, the BA.4/5 spike elicits nAbs that are highly cross-reactive for VOCs. Serological mapping studies and isolation of monoclonal antibodies from BA.4 infected individuals will provide insights into the targets of these cross-reactive neutralizing responses. These data suggest that second generation vaccines based on either Beta or Omicron BA.4 may trigger more cross-reactive nAbs than current vaccines or those based on BA.1.

## Limitations of the study

We acknowledge that the numbers of samples in each group are small. Furthermore, although we have extensive clinical follow-up, we cannot rule out the possibility that convalescent donors had experienced previous undocumented asymptomatic infection which could alter the breadth of humoral responses. Lastly, viral sequences were available only for a subset of samples in each wave, though the samples were collected when each variant dominated infections during that particular wave.

**Figure S1:**
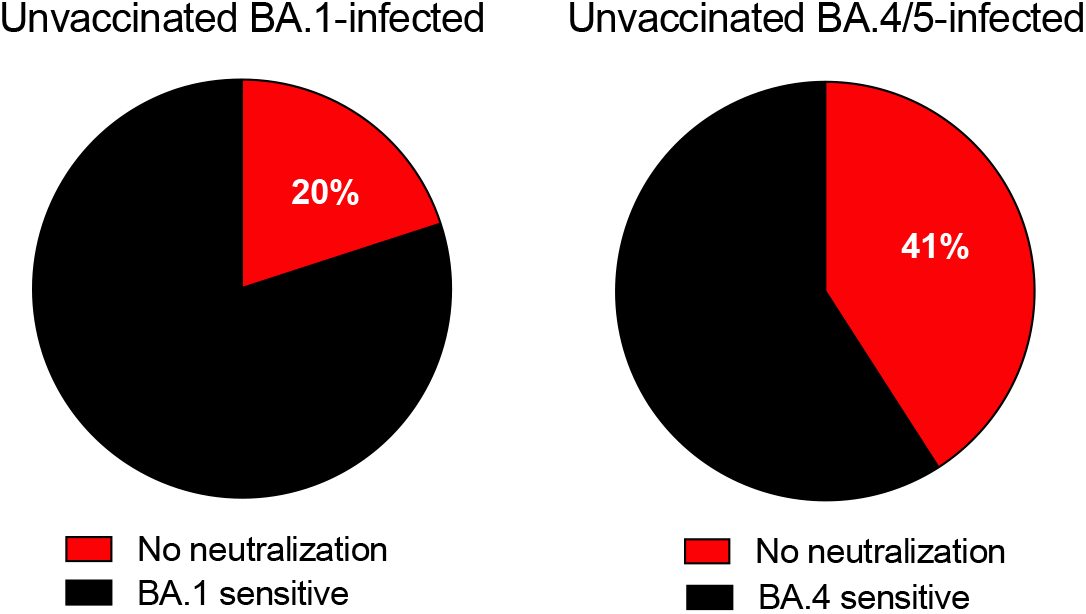
Comparison of the proportion of neutralization non-responders, defined as those individuals with no detectable neutralization to spike-matched pseudoviruses in unvaccinated convalescent donors infected with BA.1 (n=25) or BA.4/5 (n=23). Red denotes lack of neutralization, and black denotes neutralization with and ID50 of greater than 20, which is the limit of detection of the assay.

**Table S1.**
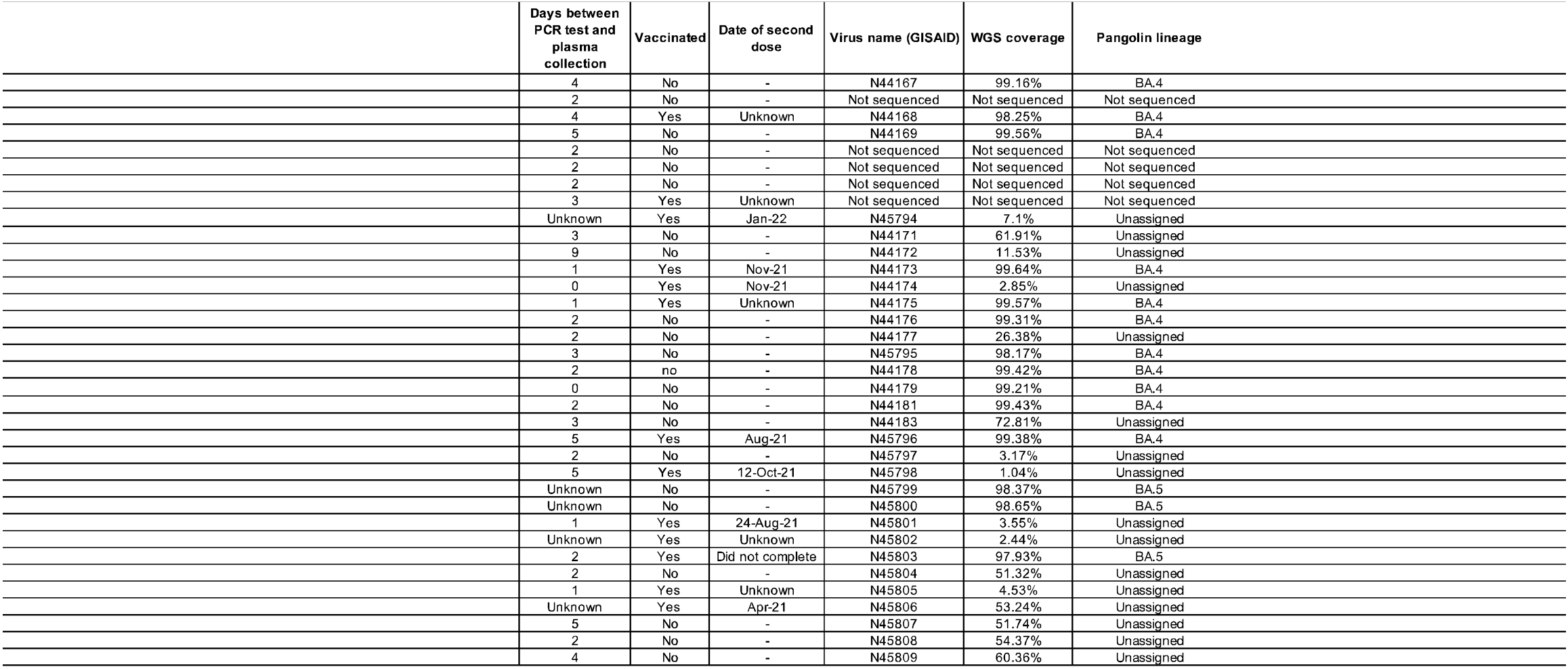
Demographic and clinical description of the cohort

## Methods

### RESOURCE AVAILABILITY

#### Lead Contact

Further information and reasonable requests for resources and reagents should be directed to and will be fulfilled by the lead contact, Penny Moore.

#### Materials availability

Materials will be made by request to Penny Moore.

#### Data and code availability

- All data reported in this paper will be shared by the lead contact upon request.
- This paper does not report original code.
- Any additional information required to reanalyse the data reported in this paper is available from the Lead Contact upon request.

### EXPERIMENTAL MODEL AND SUBJECT DETAILS

#### Human Subjects

Plasma samples from the first SARS-CoV-2 wave (D614G-infected) were obtained from a previously described cohort across various sites in South Africa prior to September 2020 (3). Second wave samples (Beta-infected) were obtained from a cohort of patients admitted to Groote Schuur Hospital, Cape Town in December 2020 - January 2021 (18). Third wave samples (Delta-infected) were obtained from the Steve Biko Academic Hospital, Tshwane from patients admitted in July 2021(15). Samples infected in the fourth COVID-19 wave of infection in South Africa were collected from participants enrolled to the Pretoria COVID-19 study cohort. Participants were admitted to Tshwane District Hospital (Pretoria, South Africa) between 25 November 2021-20 December 2021 (Table S1). In all waves, samples were collected when more than 90% of SARS-CoV-2 cases in South Africa were caused by the respective variants. Sequence confirmation was only available for a subset of samples but all the samples that were sequenced corresponded to the appropriate variant for that wave. All samples were from HIV-uninfected individuals who were above 18 years of age and provided consent. Ethical clearance was obtained for each cohort from Human Research Ethics Committees from the University of Pretoria (247/2020) and University of Cape Town (R021/2020). All patients had PCR confirmed SARS-CoV-2 infection before blood collection BTI participants were recruited from HCWs at the NICD (Human Research Ethics Committees of the University of the Witwatersrand reference number: M210465), Steve Biko Academic Hospital (Tshwane, South Africa) and Groote Schuur Hospital (Cape Town, South Africa). Written informed consent was obtained from all participants. Lack of prior infection in these individuals was confirmed by nucleocapsid ELISA.

#### Cell lines

Human embryo kidney HEK293T cells were cultured at 37°C, 5% CO_2_, in DMEM containing 10% heat-inactivated fetal bovine serum (Gibco BRL Life Technologies) and supplemented with 50 μg/ml gentamicin (Sigma). Cells were disrupted at confluence with 0.25% trypsin in 1 mM EDTA (Sigma) every 48–72 hours. HEK293T/ACE2.MF cells were maintained in the same way as HEK293T cells but were supplemented with 3 μg/ml puromycin for selection of stably transduced cells. HEK293F suspension cells were cultured in 293 Freestyle media (Gibco BRL Life Technologies) and cultured in a shaking incubator at 37°C, 5% CO_2_, 70% humidity at 125rpm maintained between 0.2 and 0.5 million cells/ml. Jurkat-Lucia™ NFAT-CD16 cells were maintained in IMDM media with 10% heat-inactivated fetal bovine serum (Gibco, Gaithersburg, MD), 1% Penicillin Streptomycin (Gibco, Gaithersburg, MD) and 10 μg/ml of Blasticidin and 100 μg/ml of Zeocin was added to the growth medium every other passage. Cells were cultured at 37°C, 5% CO_2_ in RPMI containing 10% heat-inactivated fetal bovine serum (Gibco, Gaithersburg, MD) with 1% Penicillin Streptomycin (Gibco, Gaithersburg, MD) and 2-mercaptoethanol to a final concentration of 0.05 mM and not allowed to exceed 4 × 10^5^ cells/ml to prevent differentiation.

### METHOD DETAILS

#### Spike plasmid and Lentiviral Pseudovirus Production

The SARS-CoV-2 Wuhan-1 spike, cloned into pCDNA3.1 was mutated using the QuikChange Lightning Site-Directed Mutagenesis kit (Agilent Technologies) and NEBuilder HiFi DNA Assembly Master Mix (NEB) to include D614G (original) or lineage defining mutations for Beta (L18F, D80A, D215G, 242-244del, K417N, E484K, N501Y, D614G and A701V), Delta (T19R, 156-157del, R158G, L452R, T478K, D614G, P681R and D950N), Omicron BA.1 (A67V, 69-70del, T95I, G142D, 143-145del, 211del, L212I, 214EPE, G339D, S371L, S373P, S375F, K417N, N440K, G446S, S477N, T478K, E484A, Q493R, G496S, Q498R, N501Y, Y505H, T547K, D614G, H655Y, N679K, P681H, N764K, D796Y, N856K, Q954H, N969K, L981F), Omicron BA.2 (T19I, L24S, 25-27del, G142D, V213G, G339D, S371F, S373P, S375F, T376A, D405N, R408S, K417N, N440K, S477N, T478K, E484A, Q493R, Q498R, N501Y, Y505H, D614G, H655Y, N679K, P681H, N764K, D796Y, Q954H, N969K) or Omicron BA.4 (T19I, L24S, 25-27del, 69-70del, G142D, V213G, G339D, S371F, S373P, S375F, T376A, D405N, R408S, K417N, N440K, L452R, S477N, T478K, E484A, F486V, Q498R, N501Y, Y505H, D614G, H655Y, N679K, P681H, N764K, D796Y, Q954H, N969K). Pseudotyped lentiviruses were prepared by co-transfecting HEK293T cell line with the SARS-CoV-2 ancestral variant spike (D614G), Beta, Delta, Omicron BA.1, Omicron BA.2 or Omicron BA.4 spike plasmids in conjunction with a firefly luciferase encoding lentivirus backbone (HIV-1 pNL4.luc) plasmid as previously described^7^. Culture supernatants were clarified of cells by 0.45-μM filter and stored at −70 °C.

#### Pseudovirus neutralization assay

For the neutralization assay, plasma samples were heat-inactivated and clarified by centrifugation. Heat-inactivated plasma samples from vaccine recipients were incubated with the SARS-CoV-2 pseudotyped virus for 1 hour at 37°C, 5% CO_2_. Subsequently, 1×10^4^ HEK293T cells engineered to over-express ACE-2 (293T/ACE2.MF) (kindly provided by M. Farzan (Scripps Research)) were added and incubated at 37°C, 5% CO_2_ for 72 hours upon which the luminescence of the luciferase gene was measured. Titers were calculated as the reciprocal plasma dilution (ID_50_) causing 50% reduction of relative light units. CB6 and CA1 was used as positive controls for D614G, Beta and Delta. 084-7D, a mAb targeting K417N was used as a positive control for Omicron BA.1 and Beta.

#### SARS-CoV-2 whole genome sequencing

Viral RNA was extracted using the MagNA Pure 96 DNA and Viral Nucleic Acid kit on the MagNA Pure 96 system (Roche Diagnostics) as per the manufacturer’s instructions. Quantitative PCR was performed with the TaqPath™ COVID□19 CE□IVD RT□PCR Kit (Thermo Fisher Scientific) to obtain mean cycle threshold (Ct) values for the extracts, for SARS-CoV-2 N, S and ORF1ab genes, according to the manufacturer’s instructions. The Ct values were used to split extracts into batches of n=8 per batch, with a maximum Ct range of 5 within a batch. Two batches were included per run. Extracts with <1000 copies were sequenced according to the low viral titre protocol. Batches were sequenced using the Ion AmpliSeq SARS-CoV-2 Insight Research Assay on an Ion Torrent Genexus Integrated Sequencer (Thermo Fisher Scientific). Each run completed within 24h, and included automated cDNA synthesis, library preparation, templating preparation, and sequencing. The Ion AmpliSeq SARS-CoV-2 Insight Research Assay includes human expression controls for quality control. Final SARS-CoV-2 amplicons displayed lengths ranging from 125 bp – 275 bp.

#### SARS-CoV-2 genome assemblies

Fastq files were uploaded to the SARS-CoV-2 RECoVERY pipeline (Reconstruction of Coronavirus Genomes & Rapid Analysis, v3.1) available on Galaxy ARIES (https://aries.iss.it) for genome assembly^24^. Final consensus genomes were downloaded and aligned to the reference MN908947 using NextAlign via the NextClade web portal (https://clades.nextstrain.org/, v2.0.0). Genomes were viewed in Nextclade for the presence of unknown frameshifts. Further inspection of alignments using Aliview v1.27 (https://ormbunkar.se/aliview/) revealed that these were due to single nucleotide indels and as such were likely sequencing artefacts; these were corrected to fix the open reading frames. Known frameshifts were not corrected. Nextstrain clade assignments were obtained using NextClade v2.0.0 and pangolin lineage assignments were obtained using the pangolin command line tool (pangolin v4.0.6, using the command pangolin --skip-scorpio <fasta> to avoid overwriting of BA.4 and BA.5 calls, https://github.com/cov-lineages/pangolin/issues/449). All sequences were further manually inspected to confirm lineage assignments. For all sequences that remained unassigned, clade assignments were updated to match.

### QUANTIFICATION AND STATISTICAL ANALYSIS

Analyses were performed in Prism (v9; GraphPad Software Inc, San Diego, CA, USA). Non-parametric tests were used for all comparisons. The Mann-Whitney and Wilcoxon tests were used for unmatched and paired samples, respectively. The Friedman test with Dunns correction for multiple comparisons was used for matched comparisons across variants. All correlations reported are non-parametric Spearman’s correlations. *P* values less than 0.05 were considered statistically significant.

## Acknowledgements

We thank Z van der Walt, T de Villiers, F Abdullah, P Rheeder, A Malan, W van Hougenhouck-Tulleken for clinical support. PLM is supported by the South African Research Chairs Initiative of the Department of Science and Innovation and National Research Foundation of South Africa, the SA Medical Research Council SHIP program, the Centre for the AIDS Program of Research (CAPRISA). We acknowledge funding from the Bill and Melinda Gates Foundation, through the Global Immunology and Immune Sequencing for Epidemic Response (GIISER) program. SIR is a L’Oreal/UNESCO Women in Science South Africa Young Talents awardee. The Sequencing activities for NICD (CRDM) are supported by the African Society of Laboratory Medicine (ASLM) and Africa Centers for Disease Control and Prevention through a sub-award from the Bill and Melinda Gates Foundation grant number INV-018978.

